# Gamma coherence mediates interhemispheric integration during multiple object tracking

**DOI:** 10.1101/850545

**Authors:** Nicholas S. Bland, Jason B. Mattingley, Martin V. Sale

## Abstract

Our ability to track the paths of multiple visual objects moving between the hemifields requires effective integration of information between the two cerebral hemispheres. Coherent neural oscillations in the gamma band (35–70 Hz) are hypothesised to drive this information transfer. Here we manipulated the need for interhemispheric integration using a novel multiple object tracking (MOT) task in which stimuli either moved *between* the two visual hemifields—requiring interhemispheric integration—or moved *within* separate visual hemifields. We used electroencephalography (EEG) to measure interhemispheric coherence during the task. Human observers (21 female; 20 male) were poorer at tracking objects between-versus within-hemifields, reflecting a cost of interhemispheric integration. Critically, gamma coherence was greater in trials requiring interhemispheric integration, particularly between sensors over parieto-occipital areas. In approximately half of the participants, the observed cost of integration was associated with a failure of the cerebral hemispheres to become coherent in the gamma band. Moreover, individual differences in this integration cost correlated with endogenous gamma coherence at these same sensors, though with generally opposing relationships for the real and imaginary part of coherence. The real part (capturing synchronisation with a near-zero phase-lag) benefited between-hemifield tracking; imaginary coherence was detrimental. Finally, instantaneous phase-coherence over the tracking period uniquely predicted between-hemifield tracking performance, suggesting that effective integration benefits from sustained interhemispheric synchronisation. Our results show that gamma coherence mediates interhemispheric integration during MOT, and add to a growing body of work demonstrating that coherence drives communication across cortically distributed neural networks.

## 1.1 Introduction

A highly interconnected system like the brain requires mechanisms for effective and selective neural communication. There is converging evidence that coherent (or phase-locked) neural oscillations drive this communication (the “communication through coherence” hypothesis; Fries, 2015), where networks of synchronised oscillations dynamically emerge to route information to task-relevant cortical sites (Salinas and Sejnowski, 2001; Varela et al., 2001; Buzsáki and Draguhn, 2004; Siegel et al., 2012). This concept was motivated by animal studies that demonstrated an important role for coherent gamma (35–70 Hz) oscillations within the visual systems of the cat and monkey (e.g., Womelsdorf et al., 2007; for reviews, see Fries, 2009; Uhlhaas et al., 2009). For example, Engel et al. (1991) showed that homologous spike trains recorded from cat primary visual cortex synchronised between 40 and 60 Hz. Despite the physical distance between recording sites, cortico-cortical synchronisation occurred on average with no phase-lag (i.e., a 0° offset). Moreover, sectioning the corpus callosum abolished any interhemispheric synchronisation (see also Munk et al., 1995; Kiper et al., 1999).

In humans, electroencephalography (EEG) can be used to study the role of coherent oscillations in interhemispheric integration. We devised a multiple object tracking (MOT) task that allowed us to experimentally manipulate the need for interhemispheric integration on a per-trial basis, while also allowing us to probe the relationship between interhemispheric coherence and an objective measure of integration (i.e., tracking performance; Bland et al., 2018). Our MOT arena was comprised of four quadrants, where an internal boundary manipulated the need for interhemispheric integration (**Figure 1**): objects were bound either to the two left and two right quadrants with a vertical boundary (i.e., so they remained *within* the left and right visual hemifields), or to the two upper and two lower quadrants with a horizontal boundary (i.e., so they moved freely *between* the left and right visual hemifields—requiring integration). Across a range of frequency bands chosen *a priori* (Helfrich et al., 2014), we aimed to examine the contribution of interhemispheric EEG coherence (measured between symmetrical pairs of EEG sensors) to interhemispheric integration (i.e., between-versus within-hemifield tracking) and tracking performance (i.e., both across individuals and across trials).

**Figure 1.**
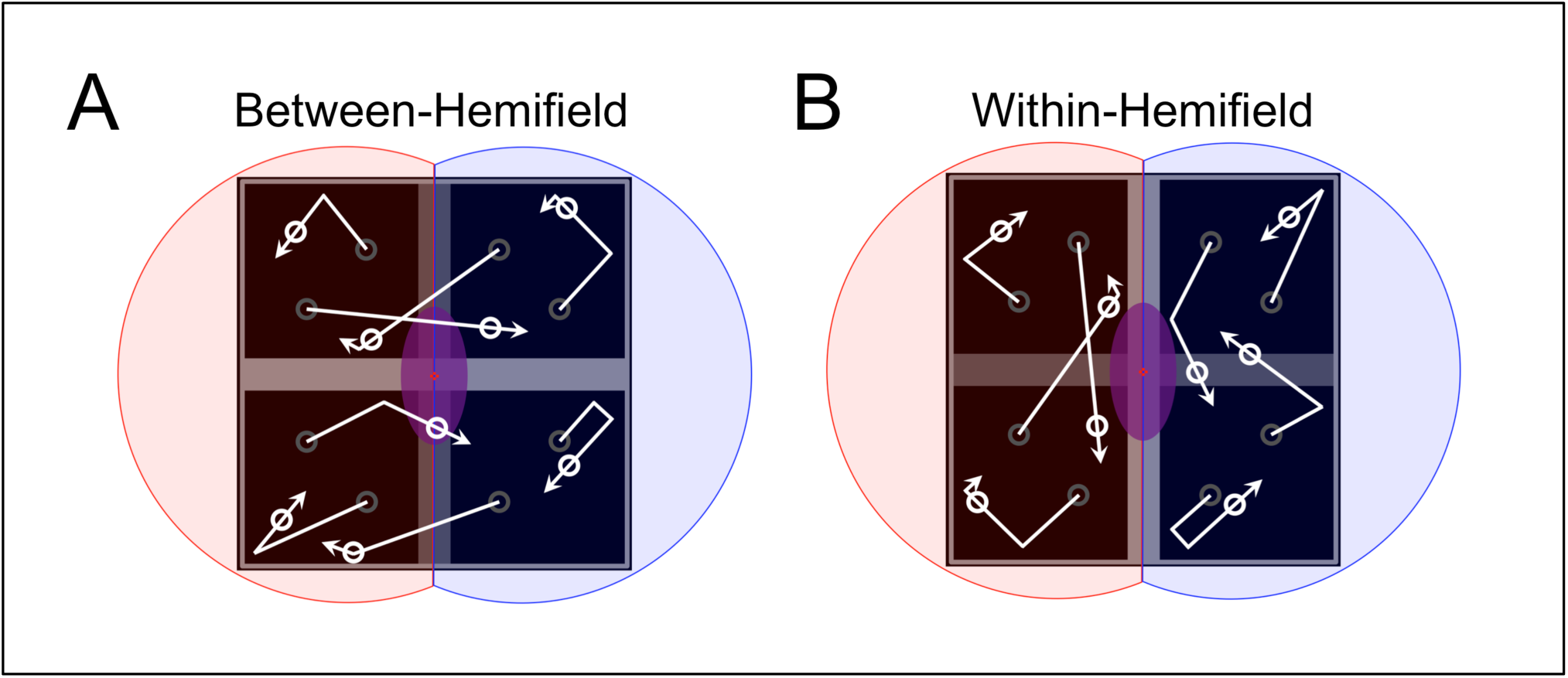
An internal boundary manipulates the need for interhemispheric integration during multiple object tracking (Bland et al., 2018). **(A)** A physical boundary oriented horizontally (with objects free to pass over the vertical boundary) restricts objects to the two uppermost and two lowermost quadrants, but allows movement between the left and right visual hemifields (red and blue shaded regions)—requiring interhemispheric integration. **(B)** A physical boundary oriented vertically (with objects free to pass over the horizontal boundary) restricts objects to the two leftmost (red) and two rightmost (blue) quadrants—never allowing movement between the left and right hemifields. Therefore, successful object tracking only requires effective and continuous interhemispheric integration during between-hemifield trials. Note that the horizontal and vertical boundaries were present during both trial types, and have been selectively darkened in this figure for emphasis only.

We expected to observe boosted coherence in the gamma band (particularly from 35–70 Hz; see Rose and Büchel, 2005; Helfrich et al., 2014; Strüber et al., 2014) when observers tracked objects *between* the visual hemifields (relative to trials in which objects remained *within* a single visual hemifield)—and primarily between EEG sensors over visual areas (i.e., parieto-occipital electrodes). We expected tracking performance to be generally worse during between-hemifield trials (reflecting a cost of interhemispheric integration; e.g., Chaudhuri and Glaser, 1991; Genç et al., 2011; Bland et al., 2018; Strong and Alvarez, 2018, 2019; Minami et al., 2019) relative to within-hemifield trials. In addition, we expected individual differences in this cost of integration to correlate with individual differences in gamma coherence (again from 35–70 Hz), since observers who perform better during between-hemifield tracking should also demonstrate greater interhemispheric coherence. Lastly, we hypothesised that gamma coherence would predict tracking performance, such that successful tracking (i.e., high-performance trials) would be more strongly associated with gamma phase-coherence than when errors were made (i.e., low-performance trials), but only for between-hemifield trials. By contrast, we predicted that within-hemifield performance would not depend on interhemispheric coherence.

## 1.2 Methods

### 1.2.1 Data

Data were collected from 40 healthy adult participants and one author (ages ranged 18–34 years; *M* = 23.46 years; 21 female, 20 male). All gave written informed consent to partake, with the protocol approved by the University of Queensland’s Medical Research Ethics Committee. No participant was excluded from analysis. Digital study materials (e.g., experiment presentation script), analysis code (e.g., behavioural analysis, EEG analysis, clustering procedure), and data (e.g., behavioural and EEG data) are archived in a publicly accessible repository (https://osf.io/sh24p/).

### 1.2.2 Task

By changing how the objects interacted with an internal boundary (**Figure 1**), objects were either restricted to separate visual hemifields (i.e., passing only over the horizontal bar and thus remaining exclusively within the left and right hemifields) or moved freely between the left and right hemifields (i.e., passing over the midline vertical bar—requiring interhemispheric integration). During each trial, participants were tasked with *covertly* tracking two or four targets among a total of eight identical objects (i.e., six or four non-targets, respectively). At the end of each trial, participants used the computer mouse to select the objects they thought were the original targets (and asked to guess if they were unsure). The trial sequence is illustrated in **Figure 2**. Across 12 blocks, participants completed equal numbers of two- and four-target trials, and between- and within-hemifield trials (16 trials pseudorandomised per block; a total of 192 trials).

**Figure 2.**
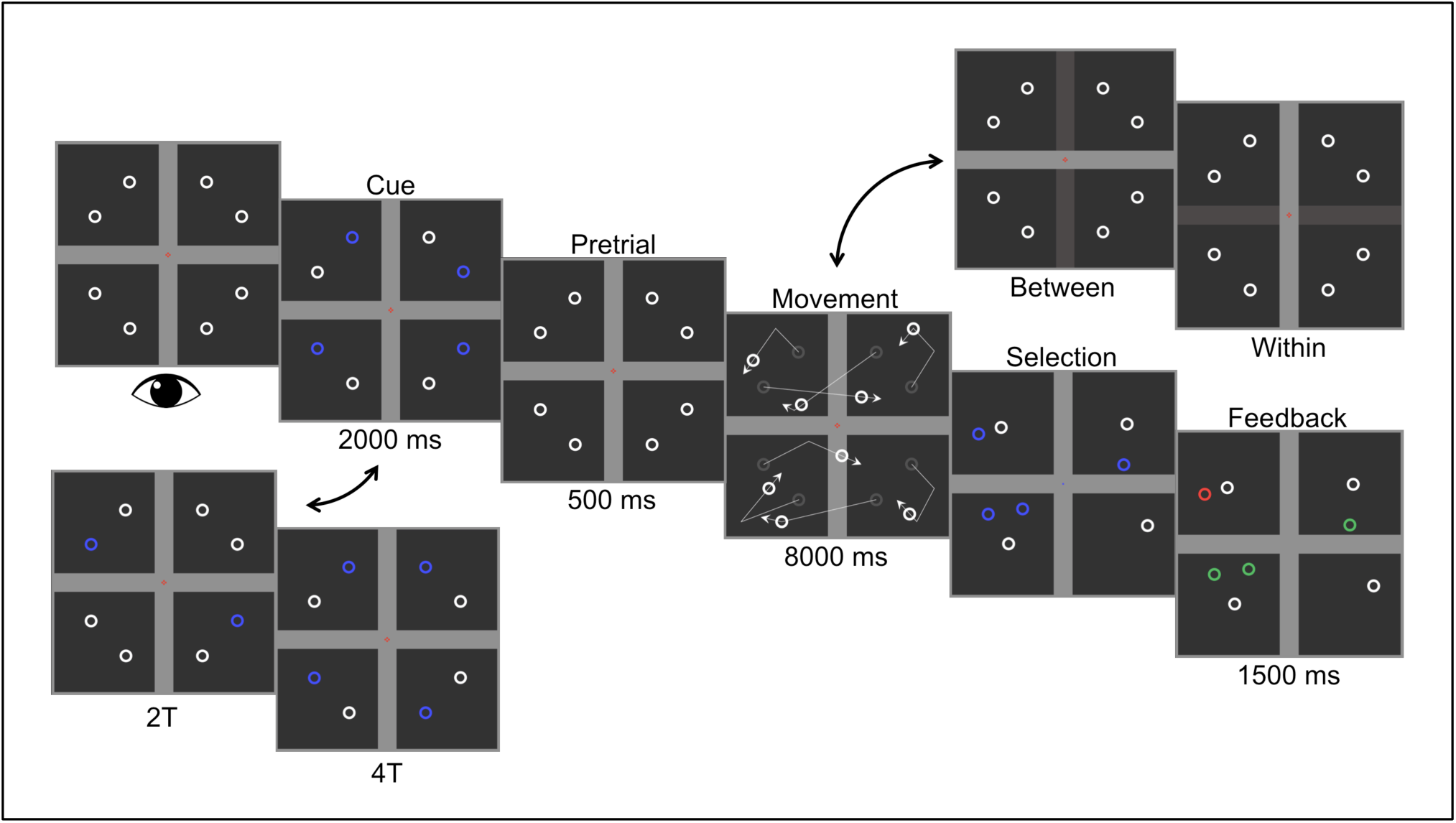
Trial sequence for the multiple object tracking task. Participants fixated centrally throughout the task. An initial cue display revealed the location of the two targets (2T) or four targets (4T) to be tracked in the upcoming trial (2000 ms; shown in blue). During the pretrial period (500 ms), all objects were indistinguishable and participants were required to remember the positions of the pre-cued targets. During the movement phase (8000 ms), objects were permitted either to pass through the vertical grey bar (for between-hemifield trials) or the horizontal grey bar (for within-hemifield trials), deflecting linearly off all other surfaces (but passing through other moving objects). Darkened grey bars (top right) illustrate the boundary through which objects could pass. Both bars were present and drawn in the same grey colour for all trials to keep trial-type ambiguous. During the selection phase, the fixation point turned into a blue cursor and participants selected with the computer mouse the objects they thought were the original targets (and asked to guess if they were unsure). Participants then received brief feedback (1500 ms) about their selections (green, correct; red, incorrect). Participants were presented trials equally divided across target number and hemifield, and pseudorandomised within each of 12 blocks (16 trials per block).

### 1.2.3 Specifications

Participants were seated 57 cm from the monitor. The MOT arena had a width and height of 24 degrees of visual angle (DVA), with horizontal and vertical bars 2.5 DVA wide (present for both trial types). The fixation cross was formed from four red squares adjacent to a central grey square (0.2 DVA square width). The circular objects had a diameter of 1 DVA, with line width 0.2 DVA. The speed of object movement (10 DVA per second) was chosen during a pilot experiment as appropriate for achieving 90% and 75% accuracy for two- and four-target trials, respectively. All objects moved linearly, with boundary deflection angles determined by the laws of reflection (i.e., angle of incidence equal to angle of reflection). It is important to note, however, that the objects passed through one another without deflecting to ensure that they would cross between adjacent quadrants multiple times per trial. Linear motion was chosen to maximise distance travelled and therefore the number of inter-quadrant crosses (∼2.64 per object over the 8 s tracking period), with the initial headings set to ensure objects would always move between adjacent quadrants (e.g., eliminating purely vertical motion during between-hemifield trials). Specifically, initial headings were sampled uniformly from ±[30–45°] and ±[135–150°] during between-hemifield trials; and from ±[45–60°] and ±[120–135°] during within-hemifield trials. During between-hemifield trials, only the horizontal bar did not allow objects to pass (**Figure 1A**); during within-hemifield trials, only the vertical bar did not allow objects to pass (**Figure 1B**). So that participants could not anticipate the type of trial, two-target trials were initialised in diagonally opposite quadrants; four-target trials were initialised with one target in each quadrant. The initial positions of all objects were kept constant across all trials, with objects beginning equidistant from fixation; their trajectories determined so that no objects overlapped at the end of the tracking period.

### 1.2.4 Eyetracking

To validate the hemifield manipulation, a subset of participants (*n* = 25) had their left eye monitored using an EyeLink 1000 (SR Research; sampled at 500 Hz). At the start of each trial, participants were required to fixate centrally (within 1 DVA of the fixation cross) before the targets were cued. The behavioural and EEG analyses were performed with trials excluded (approximately 24%) where the left eye deviated more than 1 DVA from fixation during object movement (the 8 s tracking period, **Figure 2**). The eyetracking data were also used to rule out a contribution of microsaccades to the observed coherence effects.

### 1.2.5 Electroencephalography

A custom-fit 64-channel EEG cap (EasyCap) was used, with electrodes placed in accordance with the Modified Combinatorial Nomenclature for the 10–10 system of the American Clinical Neurophysiology Society (Klem et al., 1999). Electrodes were prepared with an abrasive electrolyte gel (Abralyt, EasyCap), with impedances kept below 10 kΩ and referenced (< 5 kΩ) to the nose tip (ground at AFz; < 5 kΩ). The signals were amplified (±16 mV range) with 16-bit BrainVision hardware (BrainAmp MR Plus; Brain Products), and sampled at 1000 Hz using Recorder (BrainVision) for offline analysis in MATLAB.

All preprocessing steps and analyses were conducted in MATLAB. Filtering and independent component analysis (ICA) was performed in EEGLAB (Delorme and Makeig, 2004). The recorded EEG was zero-phase bandpass filtered from 1 Hz to 100 Hz to avoid phase distortions. The EEG was epoched and manually screened by visual inspection for unusual artefacts (e.g., jaw clenches). Fast symmetrical ICA (Hyvärinen and Oja, 2000) was then used to further remove artefacts in the EEG (e.g., blinks, microsaccades, line noise). In line with others (e.g., Hipp and Siegel, 2013; Helfrich et al., 2016b), components were removed based on three criteria: highly localised topography (e.g., line noise), abnormal power spectra (e.g., muscle activity), and high variance over trials. Importantly, these components were chosen blind to their impact on the results. Further, as the artefact from mains electricity was highly coherent across all channels and all participants, power and coherence estimates at 50 Hz were excluded from the γ_2_ band (46–70 Hz). For analysis of the phase-locking value (PLV), the EEG was further bandpass filtered over the γ_1_ band (36–45 Hz; i.e., so that only γ_1_ phase-locking was measured).

### 1.2.6 Coherence

To capture communication between the cerebral hemispheres, coherence was estimated across the 8 s tracking period between symmetrical pairs of EEG sensors. Magnitude-squared coherence (MSC) estimates were derived from Welch’s method for the cross-power spectral density and the two auto-power spectral densities, with 2 s Hamming windows with 50% overlap. The MSC between two signals *x* and *y* at frequency *f* is equal to the squared magnitude of the (complex) cross power spectral density *Pxy* of *x* and *y*, standardised by the product of the auto-spectra *Pxx* and *Pyy*. Here, *Pxx* and *Pyy* are the power estimates of *x* and *y*, and are always real-valued.

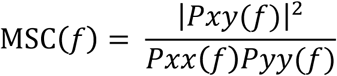

To resolve rudimentary phase information, the co-spectrum (Real) and quadrature spectrum (Imag) of the complex *Pxy* were also separately squared, giving real coherence (*Rxy*) and imaginary coherence (*Ixy*) that sum to the MSC (remembering the MSC is itself already squared). As described in **Figure 3**, the real part is largest when synchronisation occurs near 0° and 180°; the imaginary part is largest when synchronisation occurs near 90° and 270°. We chose to separately analyse the real and imaginary parts of coherence for three reasons: (1) animal and human studies suggest that coherence may occur with 0° lag between cerebral hemispheres (i.e., in the real spectrum; e.g., Engel et al., 1991; Helfrich et al., 2014); (2) the imaginary part of coherence is robust to volume conduction (e.g., Nolte et al., 2004); and (3) the real part of coherence (specifically 0° and 180°) can be targeted with rhythmic brain stimulation (e.g., Helfrich et al., 2014; Strüber et al., 2014; Bland et al., 2018).

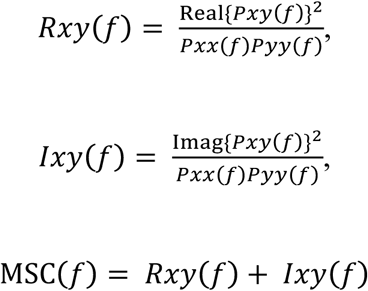

**Figure 3.**
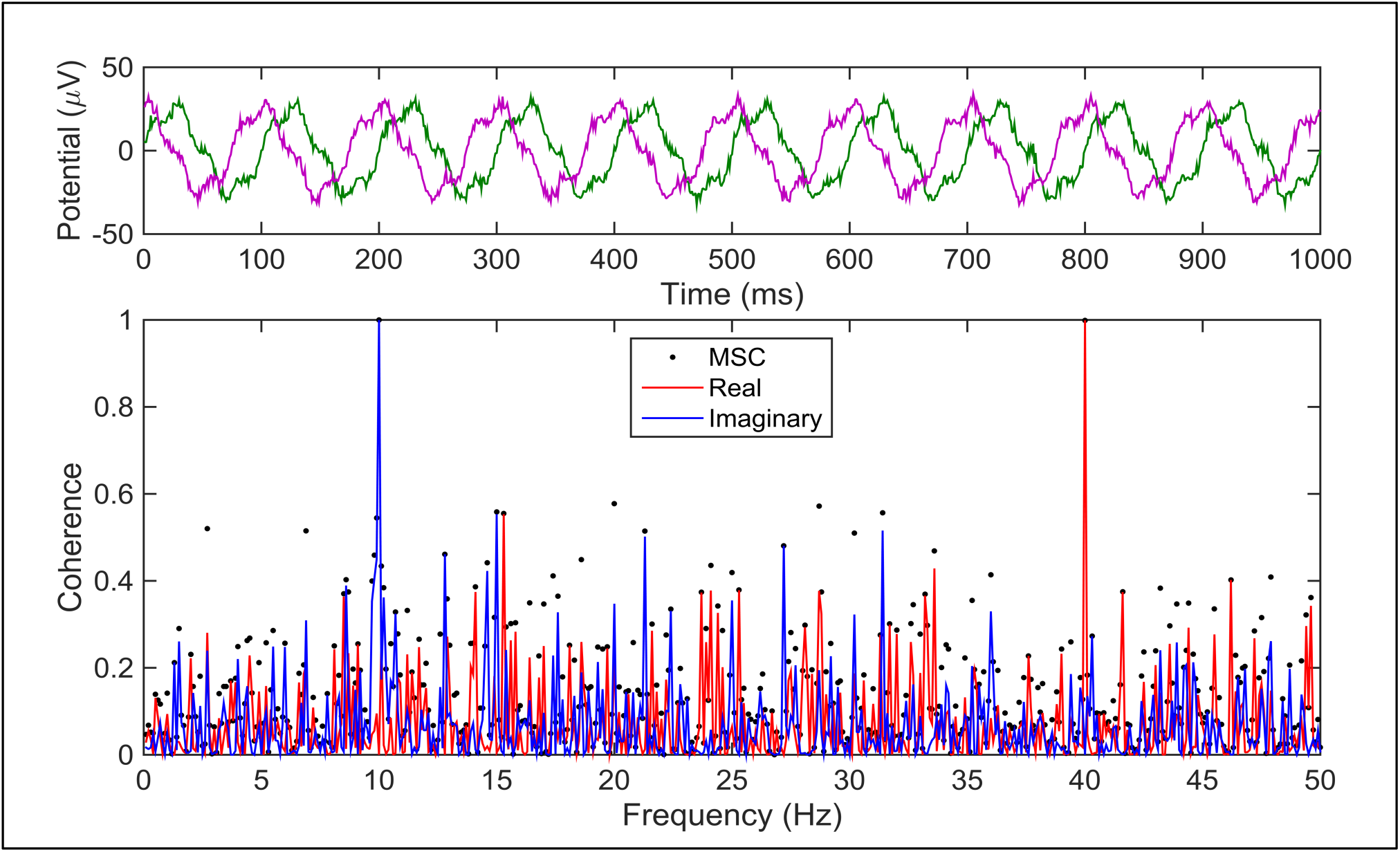
Coherence as a metric for functional connectivity in the brain. **(Top)** Two signals contain large 10 Hz and 40 Hz components, with green delayed 25 ms behind purple. This corresponds to a 90° phase-lag at 10 Hz (100 ms period) but a 0° phase-lag at 40 Hz (25 ms period). **(Bottom)** The real (red) and imaginary (blue) parts of coherence capture this phase information, with the real part largest when phase-lags are near 0° and 180°; the imaginary part is largest when phase-lags are near 90° and 270°. For all frequencies, the (squared) real and imaginary parts sum to the magnitude-squared coherence (MSC; black dots). The MSC alone does not capture any information about the ongoing phase relationships (i.e., it is approximately equal for 10 Hz and 40 Hz)—only the extent to which the two signals are linearly related. If MSC = 1, the instantaneous phase and amplitude of the two signals are linearly dependent for a given frequency (and 0 if completely independent).

Power and coherence estimates were averaged over predefined frequency bands (Helfrich et al., 2014): delta [δ: 2–4 Hz], theta [θ: 5–7 Hz], alpha [α: 8–12 Hz], low beta [β_1_: 13–25 Hz], high beta [β_2_: 26–35 Hz], low gamma [γ_1_: 36–45 Hz], mid gamma [γ_2_: 46–70 Hz, excluding 50 Hz], and high gamma [γ_3_: 71–99 Hz].

### 1.2.7 Phase-Locking Value

To compute the phase-locking value (PLV; Lachaux et al., 1999; Le Van Quyen et al., 2001) between signals *x* and *y*, we first computed their analytic signals (i.e., adding to each its Hilbert transform as the imaginary part). The instantaneous phase *ϕ* at time *t* is the four-quadrant inverse tangent of the imaginary and real components, evaluated separately for the two signals (the first and last 10% trimmed for filtering artefacts). Now, the instantaneous phase difference between signals *x* and *y* could be evaluated across the tracking period: Δ*ϕ*(*t*) = *ϕ*_*x*_(*t*) − *ϕ*_*y*_(*t*). This procedure was performed for all symmetrical sensor pairs. To evaluate the PLV at time *t* over *K* given trials:

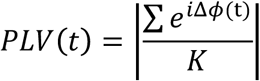

To capture differences in the strength of association between the instantaneous PLV and time, Pearson’s *r* was computed for high- and low-performance trials (**Figure 9A**). Due to the large variability in four-target tracking performance (**Figure 4A**), the criterion for “high” performance was based on individual data so that as close to equal numbers of trials were deemed “high” and “low” performance. Remembering that two-target trials were excluded from these analyses, the maximum number of targets correctly identified per trial was 4/4, with a minimum of 0/4. For example, participants with very high performance might have all 4/4 trials compared with all remaining trials (i.e., 3/4, 2/4, 1/4, and 0/4), whereas a participant with a lower overall performance might have 4/4 and 3/4 trials deemed “high” performance trials.

**Figure 4.**
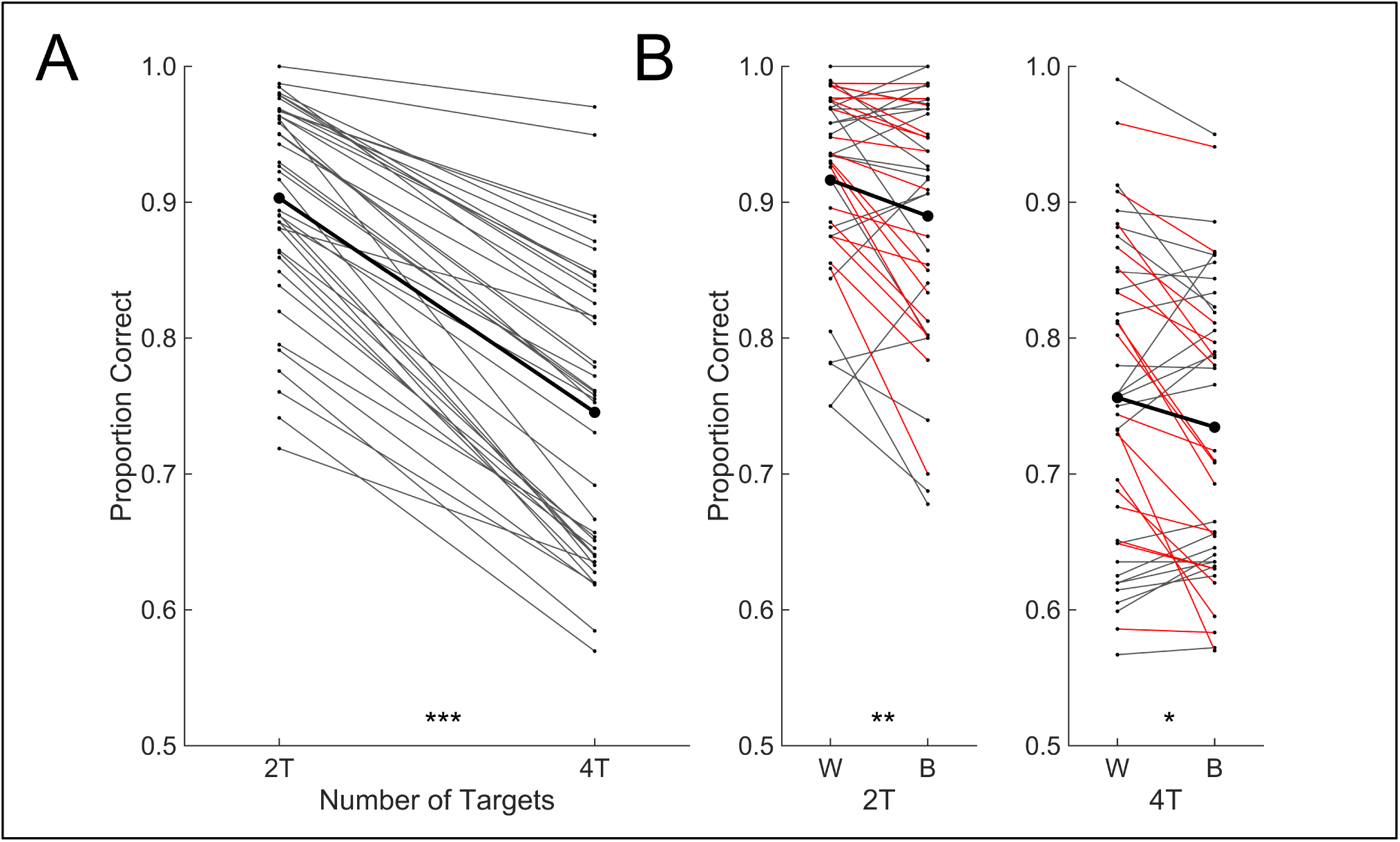
Multiple object tracking performance reflects a cost of integration. **(A)** Participants demonstrate capacity limitations, with better tracking performance in two-target (2T) trials than in four-target (4T) trials. Each individual is plotted in grey, with the mean in black. **(B)** For 2T and 4T trials, the group showed a cost of integration (i.e., worse performance for between-hemifield trials, B, versus within-hemifield trials, W). This cost of integration did not interact with the number of targets, and was largely consistent within participants (18 out of 41 participants showed a cost across both two- and four-target trials, shown in red). The remaining participants are plotted in grey. Note: *** *p* < .001, ** *p* < .01, * *p* < .05.

### 1.2.8 Cluster Permutation Analysis

All major analyses were submitted to a naïve cluster permutation algorithm that clustered the observed test statistics based on whole-scalp topographies given some criterion for clustering. To avoid spurious or sparsely connected clusters, sensors had to have at least two neighbours that also exceeded the criterion. The clustering algorithm therefore found sensors whose adjacent neighbours also exceeded the clustering criterion, only allowing sparsely connected clusters near the periphery of sensor-space. Since these were largely exploratory analyses, the clustering criterion was kept low (two-tailed α = .25 across all topographies; i.e., *t*_(40)_ = ±1.17; *r*_(39)_ = ±.17). The sensor-wise effects were permuted 10,000 times using random partitions (i.e., shuffled labels; Maris and Oostenveld, 2007). For each iteration, the permuted sensor-wise effects were summed over the observed clusters (forming distributions of the cluster statistics under the null hypothesis). From these 10,000 permuted cluster statistics, the cluster *p*–value was taken as the proportion exceeding the observed cluster test statistic (i.e., generated with the correct labels). For each family of topographies (typically 8 scalp plots), the per-cluster *p*-value was taken as the proportion of permuted cluster statistics—permuted from the maximum (largest) cluster in the family—that exceeded the observed cluster test statistic. Using the largest cluster to compute the permutation distribution allowed us to statistically control the familywise error (α = .05) across all clusters within a given family of topographies (see Maris and Oostenveld, 2007).

## 1.3 Results

### 1.3.1 Behavioural Cost of Multiple Object Tracking Between Visual Hemifields

Participants were presented with equal numbers of two- and four-target trials, and within-hemifield and between-hemifield trials (**Figure 4**). As shown in **Figure 4A**, participants were significantly better tracking two targets (*M* = 90%) than four targets (*M* = 75%), *t*_(40)_ = 16.55, *p* < .001. This difference in tracking performance supports capacity limits established in traditional MOT experiments (Scimeca and Franconeri, 2015). Consistent with previous work (Bland et al., 2018), tracking performance also reflected a cost of interhemispheric integration (**Figure 4B**): within-hemifield tracking (*M* = 84%) was significantly better than between-hemifield tracking (*M* = 81%), *t*_(40)_ = 3.64, *p* < .001. While we expected to observe a large cost of integration for four-target trials, we hypothesised this would diminish with lower demand (i.e., two-target trials). On the contrary, however, a cost of interhemispheric integration was observed for both two-target trials [*M* = −3%, *t*_(40)_ = 3.14, *p* = .003] and four-target trials [*M* = −2%, *t*_(40)_ = 2.54, *p* = .015], with no interaction, *t*_(40)_ = 0.44, *p* = .659. Unexpectedly, this cost did not correlate between two- and four-target trials, *r*_(39)_ = .21, *p* = .189, perhaps because of large individual differences in overall tracking performance: many participants with performance close to the ceiling for two-target trials would have an attenuated cost (and the same for those near the floor for four-target trials). Therefore, we tested whether those who did show a cost in two-target trials (of any size) also tended to show a cost in four-target trials. Indeed, 18 participants (**Figure 4B**, in red) showed worse between-hemifield performance for both two- and four-target trials, *P*(*n* ≥ 18 | *N* = 41, *p* = .25) = .006. Therefore, although there were large individual differences in the cost of integration, the effects were largely stable within participants.

### 1.3.2 Object Tracking Across Hemifields Modulates Interhemispheric Coherence

In line with Helfrich et al. (2014), coherence estimates were averaged over predefined frequency bands: delta [δ: 2–4 Hz], theta [θ: 5–7 Hz], alpha [α: 8–12 Hz], low beta [β_1_: 13–25 Hz], high beta [β_2_: 26–35 Hz], low gamma [γ_1_: 36–45 Hz], mid gamma [γ_2_: 46–70 Hz, excluding 50 Hz mains power], and high gamma [γ_3_: 71–99 Hz]. For interhemispheric coherence, these estimates were computed between homologous pairs of EEG sensors, resulting in symmetrical scalp topographies (**Figures 5–9**). While we had a clear prediction that the requirement for interhemispheric integration during between-hemifield tracking would modulate gamma coherence over parieto-occipital sensors, we also wanted to explore any effects over other regions. We therefore submitted whole-scalp topographies of the sensor-wise effects to a naïve clustering algorithm for permutation testing (Maris and Oostenveld, 2007). To directly test whether the need for integration modulated interhemispheric coherence, we pooled two- and four-target trials and compared the magnitude-squared coherence (MSC) observed during between-hemifield and within-hemifield trials using paired *t*-tests. We chose to pool over target number since large individual differences in tracking ability resulted in equal costs of integration for two- and four-target trials (**Figure 4B**). As shown in **Figure 5**, greater interhemispheric coherence was observed during between-hemifield tracking in almost all frequency bands (irrespective of tracking performance). As the cost of integration was largely consistent within participants (**Figure 4B**), we reasoned this was likely related to endogenous interhemispheric coherence.

**Figure 5.**
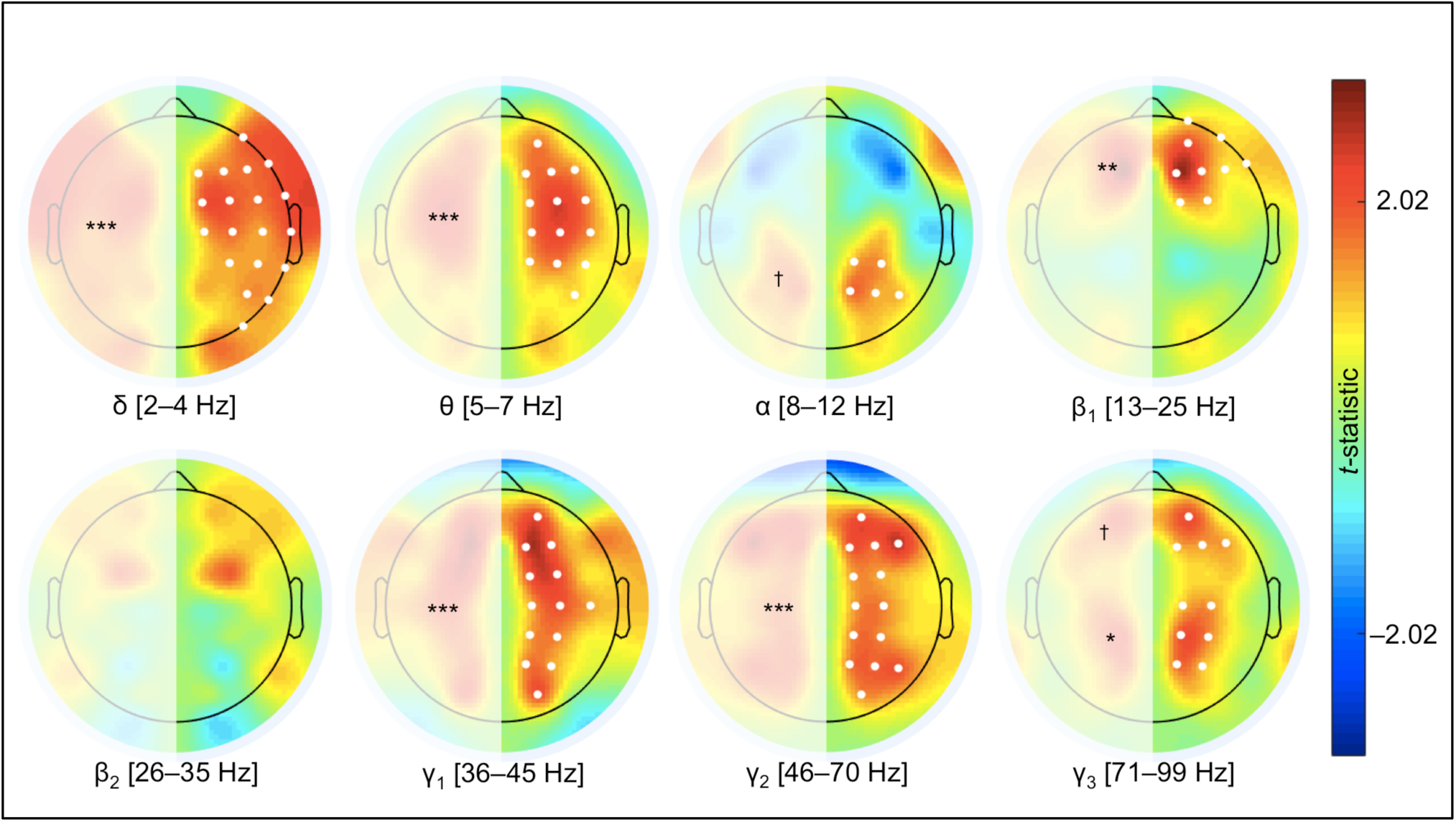
Interhemispheric coherence during between-hemifield trials versus within-hemifield trials. Scalp topographies of paired *t*-tests computed on the magnitude-squared coherence for between-hemifield versus within-hemifield trials. Positive *t*-values (in warmer colours) indicate greater coherence during between-hemifield tracking. Across almost all frequency bands, the need for integration was related to increased coherence between cerebral hemispheres, but with heterogeneous topographies. The clustered *t*-values were tested for significance using a label-shuffling permutation procedure, and the family of topographies was statistically controlled (familywise α = .05). Note: *** *p* < .001, ** *p* < .01, * *p* < .05, ^†^ failed to survive statistical correction.

We re-ran the same analysis as in **Figure 5**, but this time we split participants based on whether they showed a consistent integration cost (*N* = 18; red in **Figure 4B**) or not (*N* = 23; grey in **Figure 4B**). If between-hemifield tracking requires interhemispheric coherence, we reasoned that the cost of integration (i.e., worse between-hemifield tracking performance relative to within-hemifield trials) observed in approximately half of participants should have been driven by a failure of the two cerebral hemispheres to become coherent. Similarly, for the remaining participants with no consistent integration cost (i.e., approximately equal tracking performance for between- and within-hemifield trials), coherence should have been greater during between-hemifield tracking. Put another way, interhemispheric coherence was expected to alleviate the otherwise anticipated cost of interhemispheric integration. Consistent with this prediction, splitting the group in the manner described above revealed a strong association between the behavioural cost of integration and endogenous gamma coherence (contrast **Figure 6A** with **Figure 6B**). For the 18 participants that did show a consistent behavioural cost of integration, no changes in interhemispheric gamma coherence were observed (**Figure 6B**; i.e., the comparatively poorer between-hemifield tracking performance may be the result of a failure of the cerebral hemispheres to become coherent in the gamma band).

**Figure 6.**
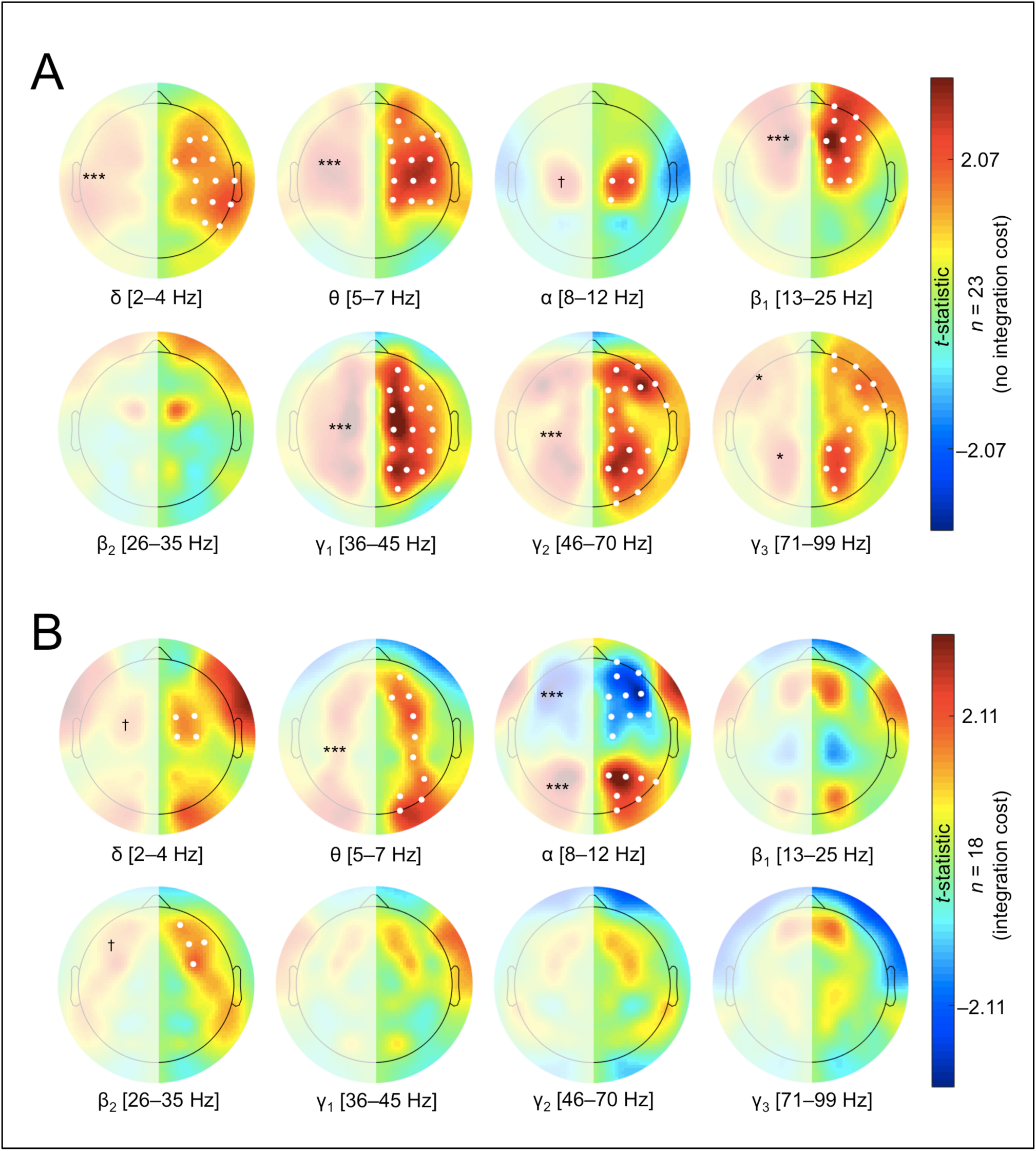
The behavioural cost of integration reflects a failure of interhemispheric coherence. Scalp topographies of paired *t*-tests computed on the magnitude-squared coherence for between-hemifield versus within-hemifield trials (as in **Figure 5**), but split by whether participants showed a behavioural cost of integration (i.e., worse tracking performance during between-hemifield trials; see **Figure 4B**). Positive *t*-values (in warmer colours) indicate greater coherence during between-hemifield tracking. **(A)** The 23 participants that did not show a consistent behavioural cost of integration (i.e., approximately equal tracking performances for between- and within-hemifield trials; grey in **Figure 4B**). For these participants, increases in interhemispheric gamma coherence were observed during between-hemifield tracking. **(B)** The 18 participants that did show a consistent behavioural cost of integration (red in **Figure 4B**). For these participants, no changes in interhemispheric gamma coherence were observed. Note: *** *p* < .001, ** *p* < .01, * *p* < .05, ^†^ failed to survive statistical correction.

Since the MSC topographies show broadband changes with the need for interhemispheric integration (**Figure 5**) and that these coherence effects depend strongly on the observed cost of integration (**Figure 6**), these results may plausibly be driven by the real or imaginary part of coherence (or both). We were therefore interested in any changes to the phasic relationships between cerebral hemispheres during between-versus within-hemifield tracking and so computed separately the topographies for the real and imaginary parts of coherence (**Figure 7**). These independent topographies were warranted because the real and imaginary parts of coherence were not strictly linearly dependent (anticorrelated) wherever a change in the MSC was observed (most frequency bands; **Figure 5**). However, since most coherence was real (especially for EEG, with high collinearity between sensors; Nolte et al., 2004), the topographies for only the real part of coherence looked very similar to the topographies for the MSC (compare **Figure 7A** to **Figure 5**).

**Figure 7.**
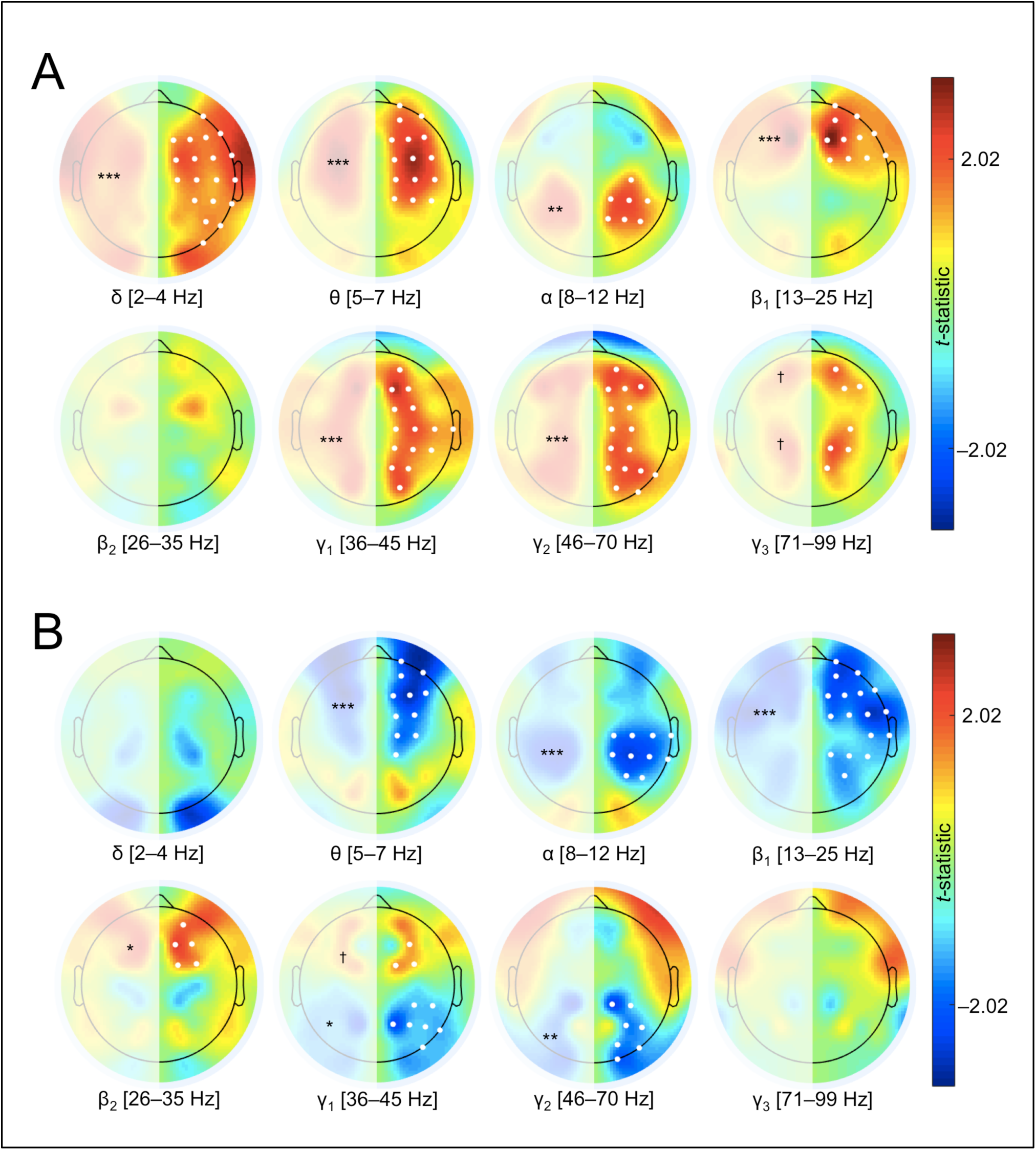
The need for integration has opposing effects on the real and imaginary parts of coherence. **(A)** Scalp topographies of paired *t*-tests computed on the real part of coherence for between-hemifield versus within-hemifield trials. Positive *t*-values (in warmer colours) indicate greater real coherence during between-hemifield tracking. Across almost all frequency bands, integration was related to increased real coherence between cerebral hemispheres, but with heterogeneous topographies. **(B)** Scalp topographies for the imaginary part of coherence for between-hemifield versus within-hemifield trials. Negative *t*-values (in cooler colours) reflect decreased imaginary coherence during between-hemifield tracking. The positive clusters over parieto-occipital sensors (α, γ_1–2_) shown in (A) were negative for the imaginary part, suggesting that interhemispheric integration requires phase alignment between the hemispheres (presumably toward a 0° offset), and actively disengages imaginary phase relationships. Note: *** *p* < .001, ** *p* < .01, * *p* < .05, ^†^ failed to survive statistical correction.

The need for interhemispheric integration modulated coherence across a range of frequencies, though with generally opposing effects for the real and imaginary parts. As shown in **Figure 7A**, between-hemifield tracking was associated with greater *real* coherence between cerebral hemispheres in almost all frequency bands, but these effects were topographically heterogeneous across frequencies. Generally speaking, frontal clusters showed boosted real coherence in the lower frequency bands (δ, θ, β_1_; *p*s < .001) during between-hemifield trials, whereas clusters containing parieto-occipital electrodes showed increases at higher frequencies (γ_1_ and γ_2_; *p*s < .001), with the exception of a posterior alpha cluster (*p* = .002). This increase in alpha coherence during between-hemifield trials was unexpected, but may explain the generally worse tracking performance observed in these trials (see also **Figure 6**). While these posterior sensors also showed high gamma coherence, the γ_1–2_ clusters were cortically distributed, suggesting that integration during MOT engages a larger network of coherent gamma oscillations than when passively observing a simple apparent motion stimulus (e.g., Helfrich et al., 2014).

As shown in **Figure 7B**, between-hemifield tracking generally disengaged imaginary phase relationships between the cerebral hemispheres. In contrast to the real part (**Figure 7A**), parieto-occipital sensors showed decreased imaginary coherence during integration in the gamma (γ_1_, *p* =.019; γ_2_, *p* = .004) and alpha (*p* < .001) bands. Similarly, the previously positive frontal clusters in the θ and β_1_ bands became negative (*ps* < .001). In fact, the only significant increase in imaginary coherence during between-hemifield tracking occurred frontally in the beta band (β_2_, *p* = .046). Together, these topographies suggest interhemispheric integration relies heavily on real phase relationships between cerebral hemispheres (presumably close to a 0° offset), while suppressing imaginary phase relationships. While several of the scalp topographies in **Figure 7** suggest that increases in the real part necessitate decreases in the imaginary part, this is only true when the MSC is unchanged (since the real and imaginary parts sum to the MSC). However, our results show that interhemispheric MSC increased during trials requiring interhemispheric integration (**Figure 5**). The observed changes in MSC could therefore be driven by either the real or imaginary part (or both), and so opposing effects would not necessarily be observed at these sensors (i.e., the effects were not linearly dependent).

### 1.3.3 Individual Differences in Coherence Predict the Cost of Interhemispheric Integration

There were large individual differences in both tracking performance (**Figure 4A**) and the cost of integration (**Figure 4B**) across participants. Since this cost of integration was largely consistent within participants (i.e., for two- and four-target trials), however, we expected individual differences in endogenous interhemispheric coherence to predict observed performance costs for between-hemifield tracking. Put another way, those better at between-hemifield tracking should tend to have more real coherence between cerebral hemispheres, particularly over parieto-occipital sensors in the gamma (γ_1–2_) band. We predicted real (and not imaginary) coherence would benefit between-hemifield tracking because, as shown in **Figure 7**, the requirement for integration up-regulates real coherence—and down-regulates imaginary coherence—at these same sensors. To directly probe the relationship between coherence and integration across participants, we computed Pearson’s *r* between the observed cost of integration (within-hemifield performance minus between-hemifield performance) and the reversed coherence effect (within-hemifield coherence minus between-hemifield coherence). Therefore, positive correlations suggest that participants who are better at between-hemifield tracking have greater interhemispheric coherence (using within-hemifield trials as a baseline for each participant). Once again, we pooled two- and four-target trials since the cost of integration did not interact with target number, and was found to be consistent within participants. This procedure was again performed separately for the real and imaginary parts of coherence, and whole-scalp topographies of *r*-values were submitted to the same naïve clustering algorithm for permutation testing (Maris and Oostenveld, 2007).

A relationship between interhemispheric coherence and the cost of integration was apparent across a range of frequencies (corroborating the between-participants approach; **Figure 6**), though with opposing effects for the real and imaginary parts. Generally speaking—from alpha through to gamma frequency bands—real coherence positively predicted between-hemifield tracking performance, but this effect moved posteriorly at higher frequencies. As shown in **Figure 8A**, real gamma coherence benefited between-hemifield tracking performance exclusively over parieto-occipital sensors (γ_1_, *p* = .009; γ_2_, *p* < .001), consistent with our hypotheses. Interestingly, real coherence at lower frequencies (θ, *p* < .001) appeared to hinder between-hemifield tracking at these same sensors. This may reflect an antagonistic relationship between gamma oscillations and lower theta–alpha oscillations (e.g., Helfrich et al., 2014, 2016a). Corroborating **Figure 7**, the relationship between integration and coherence again reversed for the imaginary part. As shown in **Figure 8B**, imaginary gamma coherence over parieto-occipital sensors was a negative predictor of between-hemifield tracking performance (γ_1–2_, *p*s < .001). Likewise, imaginary coherence at lower frequencies (δ, *p* < .001) became a positive predictor of between-hemifield performance at these same sensors. Together, these topographies (and those in **Figure 6**) suggest individual differences in endogenous interhemispheric coherence can account for some of the variability in the observed cost of integration. Those participants who were better at between-hemifield tracking showed disproportionately more real (and less imaginary) interhemispheric coherence, particularly in the gamma band and over parieto-occipital EEG sensors.

**Figure 8.**
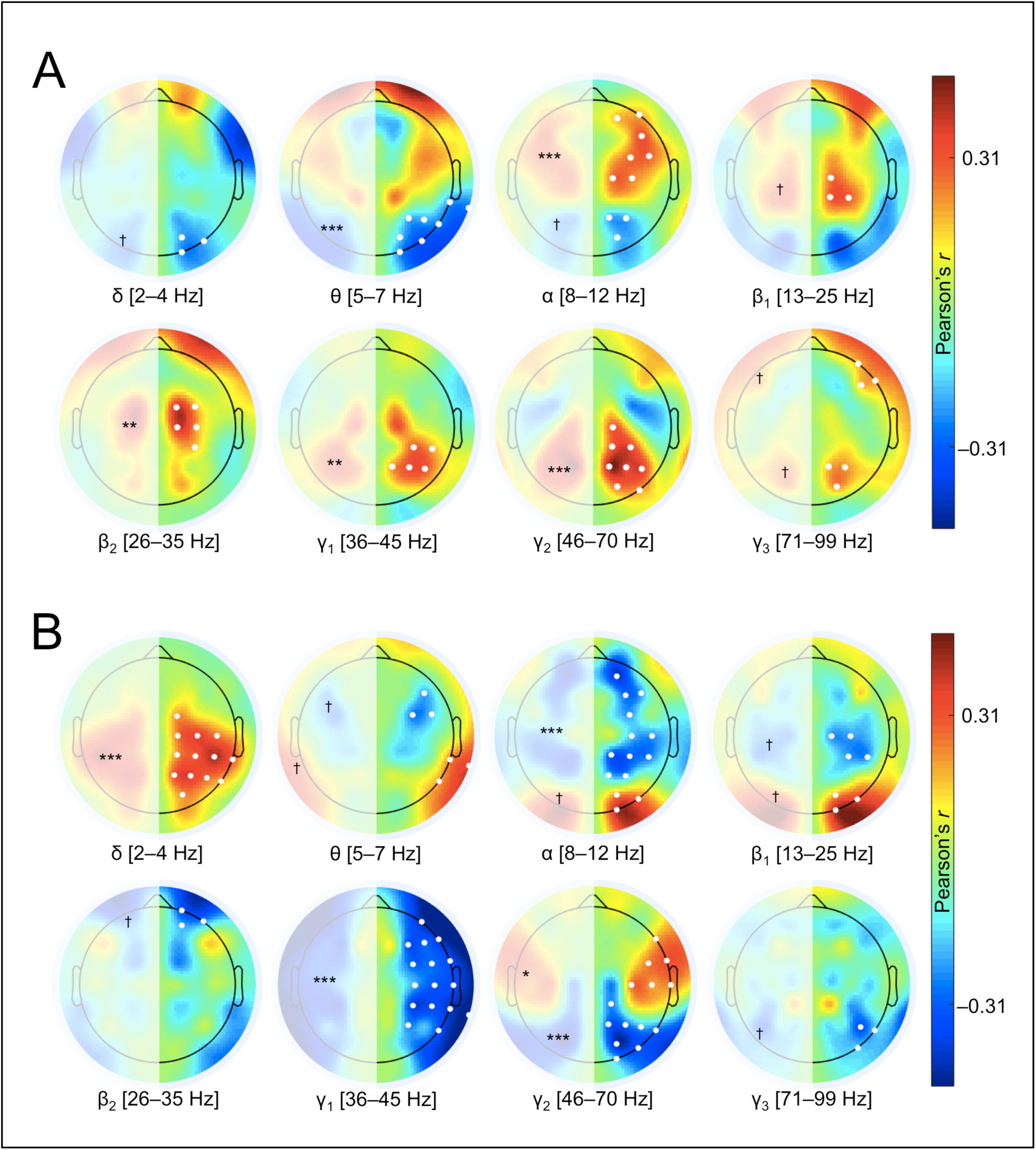
Individual differences in interhemispheric coherence predict the cost of integration, but with opposing relationships for the real and imaginary parts of coherence. **(A)** Scalp topographies of Pearson’s *r* computed between the real part of coherence (within-hemifield minus between-hemifield) and the performance cost (within-hemifield minus between-hemifield). Therefore, positive *r*-values show those with more coherence during between-hemifield trials have better between-hemifield tracking performance (using within-hemifield trials as the baseline for each participant). As predicted, real gamma coherence positively predicted between-hemifield tracking performance over parieto-occipital sensors. **(B)** Scalp topographies of Pearson’s *r* computed for the imaginary part of coherence. Negative *r*-values show those with less imaginary coherence during between-hemifield trials had better between-hemifield tracking performance. In the gamma band, imaginary coherence over parieto-occipital sensors negatively predicted between-hemifield tracking performance. Note: *** *p* < .001, ** *p* < .01, * *p* < .05, ^†^ failed to survive statistical correction.

**Figure 9.**
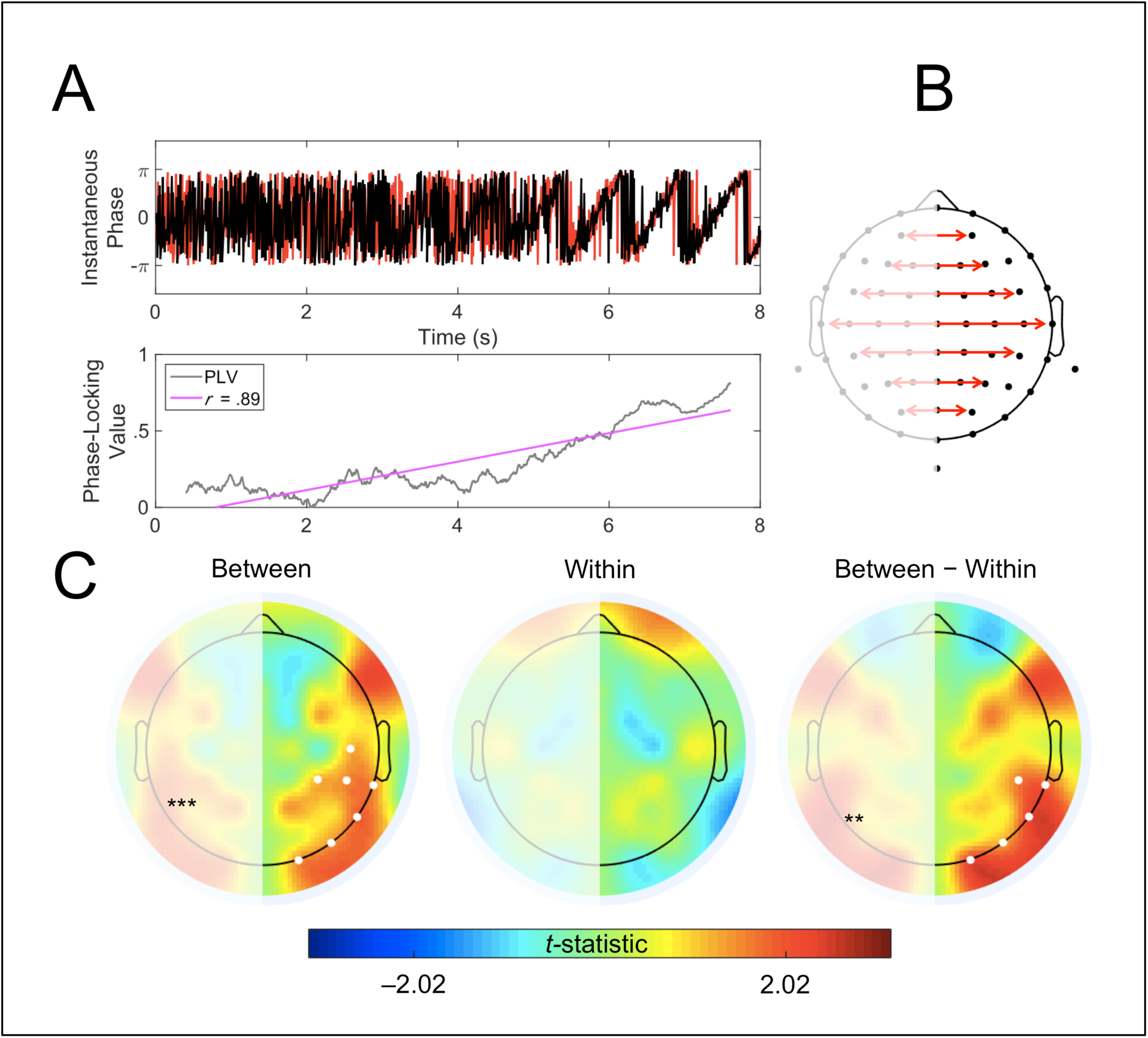
The phase-locking value discriminates high from low performance trials. **(A)** Two contrived signals (red and black; example is illustrative only) with increasingly correlated phase will have a phase-locking value (PLV) that steadily increases over time (grey). Like coherence, the PLV is bounded between 0 and 1. The strength of this relationship over time is captured using Pearson’s *r* (slope represented by pink line of best fit), where we expect there to be a stronger dependency between the gamma phase-locking value and time over the tracking period during between-hemifield trials (larger *r*-values), specifically when participants perform well versus poorly. **(B)** As for coherence topographies, the PLV was computed between symmetrical pairs of sensors. **(C)** Paired *t*-tests on the Fisher-transformed *r*-values observed during high-versus low-performance trials reveal stronger relationships during high-performance trials (positive *t*-values), but only during between-hemifield tracking—where we expect there to be a need for interhemispheric synchronisation. No difference in the strength of the correlations between high- and low-performance trials was observed for within-hemifield tracking. The interaction was significant. Note: *** *p* < .001, ** *p* < .01, * *p* < .05, ^†^ failed to survive statistical correction.

### 1.3.4 Gamma Phase-Locking Predicts Tracking Performance

Having shown that individual differences in interhemispheric coherence are associated with observed differences in the cost of integration across participants, we aimed to corroborate this result by undertaking a within-participant analysis. We predicted that trials in which participants performed well should be associated with higher gamma coherence, especially for between-hemifield trials (where integration explicitly benefits tracking). We sought to predict tracking performance from interhemispheric gamma coherence (specifically in the γ_1_ band; see Bland et al., 2018). However, since performance was near-ceiling in two-target trials (see **Figure 4A**), we analysed four-target trials only. We used a temporally resolved measure of phase coherence called the phase-locking value (PLV; Lachaux et al., 1999). This allowed us to capture changes in synchrony between homologous pairs of electrodes as before, but now at every time point over the tracking period (vastly increasing the number of observations; see **Figure 9A**). If interhemispheric integration requires reliable synchronisation between cerebral hemispheres, we would expect between-hemifield performance to benefit from a stronger dependency between the gamma PLV and time over the tracking period. By contrast, within-hemifield tracking should not depend on the strength of the association between PLV and time (i.e., for high versus low performance), since interhemispheric synchrony should not benefit performance on these trials.

To test this directly, we computed Pearson’s *r* between the PLV and time for both between-hemifield and within-hemifield trials, and split trials into *high* and *low* performance categories. Each participant therefore had four correlations computed between the gamma PLV and time: high and low performance for both between- and within-hemifield trials. We then computed paired *t*-tests on the Fisher-transformed (variance-stabilised) *r*-values for high versus low performance at each sensor, and submitted whole-scalp topographies of *t*-values to the naïve clustering algorithm for permutation testing. The EEG was zero-phase bandpass filtered in the γ_1_ band to isolate gamma phase-locked information (this frequency band was chosen *a priori*; see Helfrich et al., 2014). As shown in **Figure 9C**, the correlations between the gamma PLV and time discriminated tracking performance. Specifically, this relationship was found over parieto-occipital sensors, and only during between-hemifield tracking (*p* < .001), where phase-coherence was hypothesised to benefit performance. Explicitly, the correlations failed to discriminate high-from low-performance trials during within-hemifield tracking. This interaction was significant (*p* = .002). Together, these topographies suggest performance during between-hemifield tracking (and not within-hemifield tracking) benefits from sustained gamma synchronisation, particularly over parieto-occipital EEG sensors.

### 1.3.5 Control Analyses

Since tracking performance was generally worse for between-versus within-hemifield trials, it was plausible that the observed coherence effects reflected differences in task difficulty rather than the need for interhemispheric integration. To address this alternative explanation, we compared the MSC values in two-target between-hemifield trials with those in four-target within-hemifield trials—pitting a need for interhemispheric integration against task difficulty. These topographies are shown in **Figure 10**, and suggest the observed gamma coherence effects were driven by the need for interhemispheric integration, and not because of differences in task difficulty (or participant effort). The parieto-occipital clusters in the gamma band were the only effects that were unchanged by this control analysis (contrast **Figure 5** with **Figure 10**; i.e., boosted interhemispheric MSC during the two-target between-hemifield trials versus four-target within-hemifield trials). In corroboration of this result, **Figure 6A** also shows that the gamma coherence effects are present even in the absence of a behavioural cost of integration (i.e., between- and within-hemifield trials were essentially of equal difficulty for these individuals since they showed no cost of integration).

**Figure 10.**
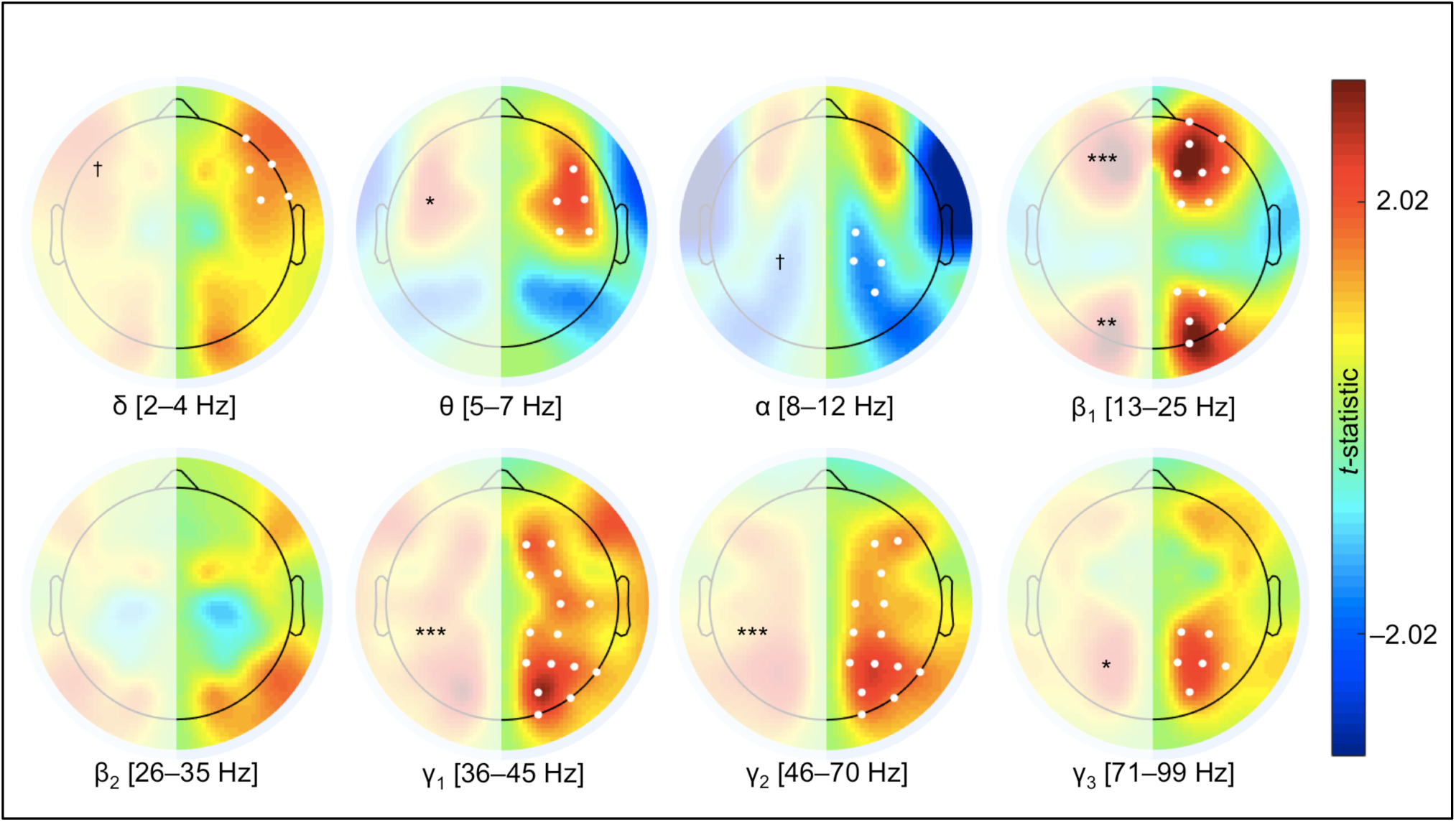
Coherence effects reflect need for integration, not task difficulty. To pit the need for integration against task difficulty, we computed paired *t*-tests on the magnitude-squared coherence observed during two-target between hemifield trials and four-target within-hemifield trials. Positive values demonstrate increased coherence during (the easier) between-hemifield trials. Note: *** *p* <.001, ** *p* < .01, * *p* < .05, ^†^ failed to survive statistical correction.

Coherence depends not only on the cross-spectrum between sensors but also on their independent auto-spectra. We therefore wanted to show the observed coherence effects occurred in the absence of any differences in EEG power (Helfrich et al., 2014). For all pre-defined frequency-bands, whole-scalp topographies were computed on the power observed during between-hemifield versus within-hemifield trials (the same auto-spectra used to compute coherence estimates in **Figure 5**). These topographies are shown in **Figure 11**, and suggest the observed gamma coherence effects were not spuriously driven by power changes. In other words, gamma oscillations were equally present during between-hemifield and within-hemifield trials, and differed only in their coherence. However, there were some highly distributed power changes in the lower frequency bands (δ, θ; *p*s < .001), and so the distributed coherence effects observed in the delta and theta bands should be considered with caution (**Figures 5–7**).

**Figure 11.**
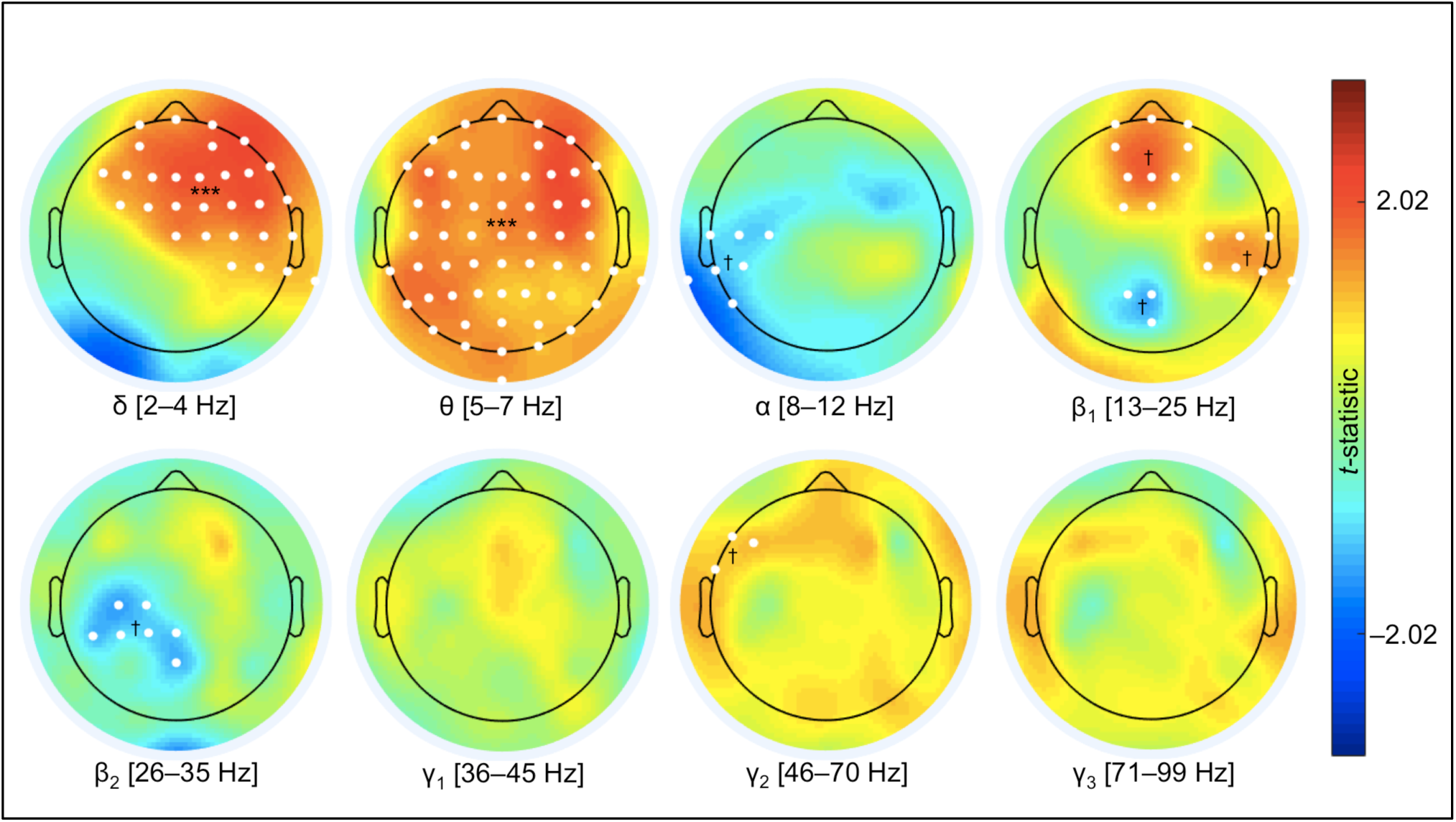
Power observed during between-hemifield tracking versus within-hemifield tracking. Positive *t*-values represent increased power for between-hemifield trials. These topographies are not symmetrical since power was computed for each sensor. Note: *** *p* < .001, ** *p* < .01, * *p* < .05, ^†^ failed to survive statistical correction.

Finally, artefacts generated by microsaccades can contaminate gamma oscillations in the EEG (Yuval–Greenberg et al., 2008). Since these artefacts are highly correlated between EEG sensors, this might also plausibly lead to spurious coherence effects, especially near 0° (i.e., the real part of coherence). Reassuringly, the topographies of the gamma coherence effects were not localised to frontal electrodes (**Figures 5–10**), and nor were there differences observed for gamma power (**Figure 11**). Nevertheless, we also analysed eyetracking data with respect to fixation and the frequency of microsaccades. A subset of participants (*n* = 25) had their left eye monitored throughout the experiment, and trials where gaze deviated more than 1 DVA from fixation (during the 8 s tracking period) were removed from all analyses (approximately 24% of all trials). We also compared the average number of microsaccades across the different trial types. In line with our other experiment using this task (Bland et al., 2018), this revealed no effect of hemifield [*t*_(24)_ = 0.62, *p* = .539] or target number [*t*_(24)_ = 1.34, *p* = .194], suggesting the observed coherence effects were not confounded by microsaccades. The whole-scalp topographies were very similar with and without the inclusion of participants without eyetracking (*n* = 16).

## 1.4 Discussion

Our findings suggest an important role for coherent gamma oscillations (∼35–70 Hz) in interhemispheric integration, as assessed in a visual multiple object tracking (MOT) task with concurrent electroencephalography (EEG). We found that trials requiring integration were associated with increased interhemispheric coherence (**Figure 5**). We exploited observed differences in the cost of integration—specifically, poorer performance during between-hemifield trials relative to within-hemifield trials (**Figure 4B**)—to show that this difference was associated with individual differences in endogenous gamma coherence (**Figures 6 and 8**). By splitting the magnitude-squared coherence (MSC), we show that between-hemifield tracking (and tracking performance) was associated with increased real gamma coherence (with phase-lags near 0° and 180°; **Figure 7A**), with the opposite observed for imaginary phase-relationships (phase-lags near 90° and 270°; **Figure 7B**). We also found, using a time-continuous measure of phase coherence (phase-locking value, PLV), that tracking performance was predicted by gamma synchrony [36–45 Hz] over time, but only during trials in which interhemispheric coherence should benefit performance; namely, during between-hemifield trials (**Figure 9**).

### 1.4.1 Real versus Imaginary Coherence

The magnitude-squared coherence (MSC, **Figure 3**) provides no information about the phase-lag between coherent oscillators. Since the real part of coherence dominates the EEG (i.e., many sensors detect the same source activity; ‘volume conduction’ at 0°), some investigators have looked at just the imaginary part of coherence (e.g., Nolte et al., 2004; Helfrich et al., 2016b), but at the expense of ignoring true neural interactions with a 0° offset. The real part still provides valuable information about functional cooperation, especially because distributed networks can oscillate with no phase-lag (e.g., Frien et al., 1994; Roelfsema et al., 1997; Rodriguez et al., 1999; Fell et al., 2001). However, even though zero-lag coherence can signify important physiological mechanisms, many factors—like electromyographic artefacts—can contribute spuriously to such observations (Buzsáki and Schomburg, 2015). In our experiment, within-hemifield trials acted as a rigorous baseline for between-hemifield trials: the two trial types were otherwise identical in every respect, and differed only in whether objects could move between the left and right visual fields. Therefore, observed coherence effects are not easily attributable to volume conduction or a common reference (Helfrich et al., 2014), since these should not change with trial type.

Generally speaking, **Figures 7 and 8** illustrate typically opposing relationships between the real and imaginary parts of coherence and interhemispheric integration. For example, the need for integration was associated with increases in the real spectrum (**Figure 7A**), but decreases in the imaginary spectrum (**Figure 7B**). Over parieto-occipital EEG sensors, this appeared particularly true in the γ_1–2_ bands (36–70 Hz), suggesting interhemispheric integration not only requires gamma synchrony, but actively down-regulates imaginary coherence (presumably reflecting a phase-shift toward 0° synchrony). Similarly, real (but not imaginary) gamma coherence positively predicted between-hemifield tracking across participants at these same sensors (**Figure 8**).

### 1.4.2 Communication Through (Gamma) Coherence

The communication through coherence hypothesis (Fries, 2015) predicts a need for synchronised (phase-locked) neural oscillations for effective and selective inter-regional brain communication. An earlier version of the communication through coherence hypothesis posited that gamma coherence was important for highly localised communication; coherence at lower frequencies might therefore foster longer-range communication (e.g., von Stein and Sarnthein, 2000) so as to keep phase-synchronisation at a near-zero offset despite increasing cortical distance (Fries, 2005). However, more recent work has shown that long-range synchronisation still occurs in the gamma band (e.g., Gregoriou et al., 2009; Bosman et al., 2012; Baldauf and Desimone, 2014; Bastos et al., 2015a), but increasingly non-zero phase-lags often emerge because of conduction delays (reviewed in Bastos et al., 2015b). By contrast, we show here that interhemispheric integration in human observers appears to benefit from synchrony with no phase-lag, corroborating other interhemispheric integration experiments (e.g., Engel et al., 1991; Helfrich et al., 2014).

It is not obvious why interhemispheric integration opposes this trend (i.e., interhemispheric coherence tends not to be imaginary, though see Helfrich et al., 2016b). A trivial case for a 0° phase-lag between distant oscillators is when the conduction delay is equal to the oscillatory period, but the observed increase in real coherence in our experiment was cortically distributed, and was spread across multiple frequency bands. Vicente et al. (2008) demonstrated that two neuronal groups that are bidirectionally connected to a third population can display coherence with no phase-lag, and many modeling studies have shown that zero-lag synchrony is biologically plausible (e.g., Chawla et al., 2001; Rajagovindan and Ding, 2008), even in the gamma band (Traub et al., 1996). These models highlight the need for recurrent connectivity and positively correlated input to achieve network synchrony with a 0° offset (see Bastos et al., 2015b). The symmetrical, bidirectional communication that occurs during interhemispheric integration may therefore exhibit a net 0° offset, despite unidirectionally non-zero transcallosal conduction delays.

### 1.4.3 Entraining Coherent Gamma Oscillations

There is a long history to the idea that coherent gamma oscillations mediate interhemispheric integration. Despite early evidence that gamma synchrony occurs with no phase-lag between cerebral hemispheres (e.g., Engel et al, 1991), the nature of this phase relationship in human studies has been largely overlooked, in large part because of the typical measures used to quantify functional connectivity in the EEG (e.g., the magnitude-squared coherence, MSC; **Figure 3**). Nevertheless, some evidence that gamma synchrony occurs with a 0° offset during interhemispheric integration comes from work employing rhythmic brain stimulation. Coherent neural oscillations may be entrained using transcranial alternating current stimulation (tACS), which allows experimental control over the ongoing phase relationship between cerebral hemispheres when applied bilaterally (see Bland and Sale, 2019). Helfrich et al. (2014) demonstrated that 40 Hz tACS applied in-phase (0° offset) biased perception of an ambiguous apparent motion stimulus toward the horizontal (which requires interhemispheric integration since the tokens move between visual hemifields). In-phase tACS also boosted endogenous gamma MSC, demonstrating a causal role for it in integration. In support of this, we demonstrate here that integration in an object tracking task also benefits from real interhemispheric gamma coherence, presumably with the same 0° offset (though see Bland et al., 2018).

### 1.4.4 Coherence and Multiple Object Tracking

In the classical multiple object tracking (MOT) paradigm, a subset of visually identical objects must be tracked over an extended period of time (typically 8–10 seconds), ignoring all irrelevant nontargets. As each object moves independently and continuously across the visual field, successfully identifying the targets at the end of this period requires the sustained, multifocal distribution of attention (Cavanagh and Alvarez, 2005). While dozens of behavioural experiments have assessed the limits of MOT and the various parameters that influence performance (e.g., object speed, number of targets; for a review, see Scimeca and Franconeri, 2015), few studies have provided a mechanism underlying MOT in terms of the neural correlates of tracking performance. During MOT, attentional resources have been shown to be largely independent between visual hemifields (e.g., Alvarez and Cavanagh, 2005; Störmer et al., 2014), and our results are consistent with this account. Since between-hemifield tracking uniquely requires objects be transferred between these independent resources (i.e., there is a need for interhemispheric integration), a failure of the cerebral hemispheres to become coherent (as observed in approximately half of the participants) may cause the observed decrease in between-hemifield tracking performance (i.e., an interhemispheric “crossover cost;” e.g., Chaudhuri and Glaser, 1991; Genç et al., 2011; Bland et al., 2018; Strong and Alvarez, 2018, 2019; Minami et al., 2019).

Evidence from neuroimaging studies suggests MOT engages a distributed, bilateral cortical network (Howe et al., 2009; Jahn et al., 2012; Alnæs et al., 2015), and that effective connectivity across this network is presumed to foster successful tracking (at least when objects cross between visual hemifields; see also Minami et al., 2019). Adding to this literature, here we find that coherence effects are maximal over parieto-occipital EEG sensors and in the gamma band when participants track visual objects across the left and right visual fields. While previous research has had some success in establishing neural correlates of hemifield crossover (Minami et al., 2019), different tracking strategies (e.g., Merkel et al., 2014, 2015), attentional load (e.g., Sternshein et al., 2011), and sustained attention (Drew and Vogel, 2008), explicit attempts to predict tracking performance have been largely unsuccessful (e.g., predicting reaction time but not accuracy; Störmer et al., 2013). Here we show that gamma coherence can predict tracking performance both across participants and across trials, and propose real coherence as a plausible neurobiological marker for effective interhemispheric integration and associated between-hemifield tracking.

### 1.4.5 Conclusion

How information is routed between task-relevant cortical regions is only rudimentarily understood, with most accounts emphasising the need for coherent (synchronised) neural oscillations. Tasks requiring interhemispheric integration provide a tool to assess electrophysiological correlates of functional cooperation (i.e., between cerebral hemispheres). By using multiple object tracking, we were able to manipulate the need for interhemispheric integration on a per-trial basis, while also having an objective measure of integration efficacy (i.e., tracking performance). We show that tracking performance reflects a cost of integration, which correlates with individual differences in interhemispheric coherence. Gamma coherence appears to uniquely benefit between-hemifield tracking, predicting performance both across participants and across trials. We have demonstrated that real (but not imaginary) interhemispheric coherence is not only related to the need for integration, but also accounts for observed differences in the cost of interhemispheric integration. These findings support the communication through coherence hypothesis, and corroborate a growing literature that suggests that coherent gamma oscillations can foster communication over cortically distributed networks.

